# Wnt-7a-positive dendritic cytonemes induce synaptogenesis in cortical neurons

**DOI:** 10.1101/2023.02.17.528927

**Authors:** Thomas M. Piers, Seema C. Namboori, Akshay Bhinge, Richard Killick, Steffen Scholpp

## Abstract

**Summary:** Neuronal circuits evolve as a precisely patterned network. In this context, a growing neuron must locate the appropriate target area on a neurite of a neighbouring cell with which to connect. Controlled target selection involves dendritic filopodial contacts and requires the exact apposition of synaptic components. Calcium signalling has been postulated to trigger the transformation from dendritic filopodia into functional synapses. However, calcium is a rather unspecific signalling system, and it needs to be clarified how the exact development of synaptic connections is controlled. Similarly, Wnt/β-catenin signalling promotes synapse formation; however, how secreted Wnts induce and maintain synapses on neuronal dendrites is not well understood. Here, we show that Wnt-7a is tethered to the tips of dynamic dendritic filopodia during spine formation in human cortical neurons. These filopodia can activate Wnt signalling precisely at the contact sites on the dendrites of an adjacent neuron. Subsequently, local calcium transients can be observed at these Wnt-positive contact sites. Depleting either the filopodial-loaded Wnt or the extracellular calcium pool blocks the clustering of pre- and post-synaptic markers, hence the establishment of stable connections. Therefore, we postulate that local Wnt-7a signalling from the tip of the dendritic filopodia, verified by simultaneous calcium signalling, provides an elegant mechanism for orchestrating focal synapse maturation.

## Introduction

The formation of synaptic connections, or synaptogenesis, is fundamental to developing a functional nervous system and expressing complex sentient behaviours, including learning, emotion, memory formation, and adaption. Synaptic formation requires precise apposition of pre- and postsynaptic components, and connections, whilst perceived to be stochastic, rarely form on the first cue, suggesting the involvement of dynamic, highly regulated dendritic filopodia that make contact and initiate signalling^1^.

Substantial progress has been made in identifying molecular processes involved in synaptic formation. In particular, cell adhesion molecules (CAMs) play a crucial role in synaptogenesis, including L1-CAM and Contactin at the presynaptic side and Neuroligins and N-Cadherin at the post-synaptic side^2,3^. However, a molecular understanding of how synaptic connections are initiated between neurons needs to be better understood. For example, what causes contacts at specific sites to differentiate into synapses but not others? How do pre- and post-synaptic neurons recognise each other and establish a stable connection as an excitatory synapse?

Secreted signalling factors are crucial for promoting synaptogenesis^4,5^. Among these growth factors, the Wnt signals play a prominent role in orchestrating adult neural development and neuronal plasticity^6^. Strong evidence suggests that Wnt proteins not only influence cellular behaviour at the tissue level but can also activate localised signalling at specific sites on the plasma membrane that regulate dynamic cytoskeletal changes^7,8^. More specifically, the Wnt/β-catenin signalling pathway triggered by Wnt7a has been shown to activate an essential cascade required for promoting the transformation of spines into a mushroom-like morphology, which can be seen as an indicator for initiating and forming synapses^9,10^. However, precisely how Wnt7a signalling induces and maintains synapses remains controversial.

Historically, the involvement of Wnt signalling in synaptogenesis has focussed on a diffusion hypothesis, whereby Wnt ligands are released from postsynaptic cells acting in a retrograde fashion to recruit presynaptic proteins, followed by bidirectional Wnt signalling and subsequent dendritic spine morphogenesis^11^. However, two issues with this model are the unfocussed nature of diffusion and the hydrophobicity of Wnt ligands. Therefore, how these ligands are targeted to specific regions on receiving neurons, and how these lipophilic growth factors are trafficked intercellularly through the aqueous extracellular space is unclear. Recently, we and others have identified filopodia, also known as cytonemes, as an essential transport mechanism. Cytonemes can bring lipophilic Wnt proteins to receptors on a target cell without an extracellular release from the donor cell^12–15^. Furthermore, recent studies in *Drosophila* have shown that these cytoneme contacts have structural similarities with neuronal synapses and that signalling facilitates glutamatergic signalling in this context^16^. Collectively, these findings have started to alter our mechanistic understanding of Wnt signalling fundamentally. Based on these data, we propose that dendritic filopodia are characteristically similar to Wnt signalling cytonemes and that their contact sites are precursors to nascent synapses.

Using super-resolution lattice structured illumination microscopy (LSIM) microscopy and mature human iPSC-derived cortical neuron cultures, we show that Wnt-7a-positive filopodia contact opposing neurons leading to clustering of the Wnt co-receptor LRP6. Consequently, we find calcium signalling at these contacts. Following local Wnt/β-catenin and calcium signalling, we observe the clustering of PSD95 on the dendritic filopodia membrane and Bassoon on the opposing, pre-synaptic side followed by a morphological transformation from dendritic filopodia into mushroom-like spines. Tethering Wnt-7a to the dendritic filopodia plasma membrane does not alter this process. Consistently, the inactivation of specifically membrane-tethered Wnts inhibits synapse formation. Thus, we conclude that dendritic filopodia resemble many features of Wnt-bearing cytonemes. Therefore, the transport of Wnt-7a by dendritic filopodia is a crucial prerequisite in forming excitatory synapses at a precise location and establishing an accurate spine pattern of the cortical neuronal network.

## Results

### Most dendritic protrusions harbour Wnt-7a and induce clustering of LRP6

To map the location of Wnt-7a during synaptogenesis, we analysed dendritic protrusions in iPSC-derived human cortical neurons (iPS-N) at 60 days in vitro (60DIV) using super-resolution LSIM with a lateral resolution of 60nm. We found the Wnt ligand, Wnt-7a, present on almost 74% of all protrusions (Figures 1A, B and S1). Of this majority of Wnt-7a-positive protrusions, co-localisation with clusters of the Wnt/β-catenin pathway coreceptor LRP6 in apposed neurites correlated with the maturity of the protrusion, suggestive of trans-activation of this pathway from the post-synaptic neuron to the pre-synaptic side (Figures 1A and C). Signalling filopodia harbour specific proteins required for the trafficking of Wnt proteins^13,17^. Similarly, we found co-localisation of Wnt-7a with the membrane-bound scaffolding protein and Wnt cytoneme marker Flotillin-2 (Flot-2) and the Wnt ligand transporter, Wntless/evenness interrupted (Wls/Evi) (Figures 1D-F and H-K). Interestingly, the actin motor Myosin 10 (Myo-10) that regulates the formation and elongation of filopodia did not co-localise on Wnt-7a-positive dendritic protrusions (Figure 1D).

**Figure 1.**
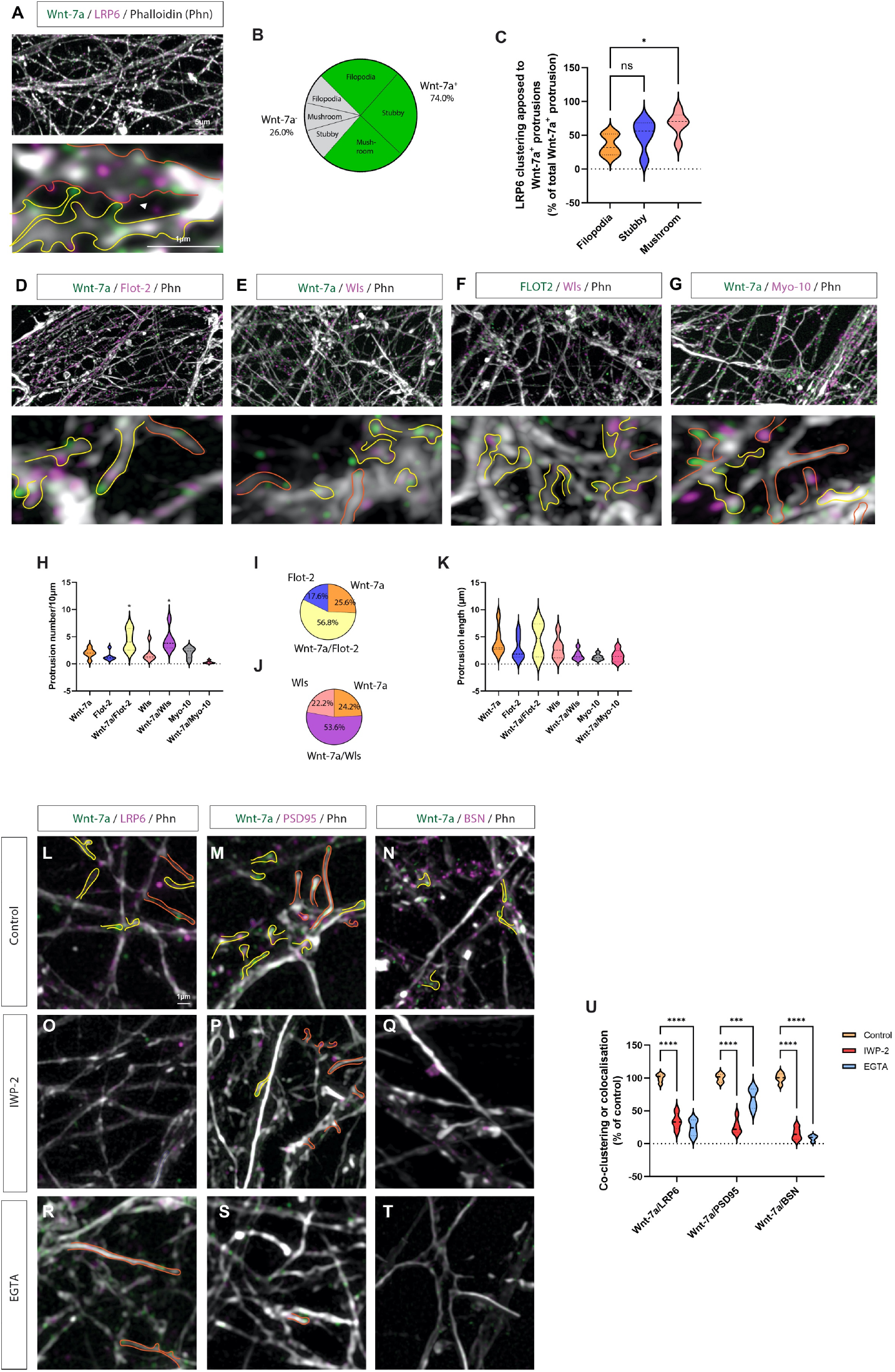
Characterisation of dendritic filopodia as Wnt-signalling filopodia. (**A**) Co-staining of iPSC-derived cortical neurons (DIV60) to observe Wnt-7a and LRP6 localisation. Wnt-7a can be seen localised on dendritic protrusions (yellow outlines), whereas LRP6 generally localises at apposing membranes (orange outlines). Wnt-7a-positive protrusions also harbour proteins associated with Wnt-signalling filopodia, such as Flotillin-2 (Flot-2; **B**; yellow outlines) and Wntless/evenness interrupted (Wls’; **C**; yellow outlines), whereas Wnt-signalling filopodia marker Myo-10 localise to a Wnt-7a-negative subset of protrusions (**D**; orange outlines). Flot-2 and Wls also co-colocalise on the filopodia (**E**; yellow outlines). (**F**) Quantification identified a subset of Wnt-7a-positive protrusions, with a significant difference observed in the length of Wnt-7a-positive filopodia, compared to Wnt-7a-negative filopodia. (**G**) Quantification of co-clustering of LRP6 and Wnt-7a at apposing membranes identified a significant increase in co-clustering of Wnt-7a-positive mushroom-shaped protrusions and LRP6 when compared to co-clustering at filopodia contacts. (**H**) Quantification of protrusion number based on cytoneme markers found significantly more Wnt-7a/Flot-2-positive and Wnt-7a/Wls-positive protrusions when compared to Wnt-7a-only positive protrusions, and over 75% of all protrusion analysed were Wnt-7a positive (**I & J**). (**K**) Quantification of protrusion length found no difference based on cytoneme protein expression (**K**). The co-clustering of Wnt-7a-positive protrusions and apposed LRP6 (**L**) was dependent on both palmitoleation (**O**) and calcium signals (**R**). Furthermore, colocalisation of Wnt-7a with the post-synaptic protein PSD95 (**M, P, S**) and the pre-synaptic protein Bassoon (BSN; **N, Q, T**), was also dependent on palmitoleation and calcium signalling, as quantified in (**U**). Statistical significance was addressed using one-way ANOVA with Dunnett’s multiple comparison test to compare relevant controls within groups. *P < 0.05; **P < 0.01; ***P < 0.005; ****P < 0.001.

We further found that the clustering of LRP6 in neurites apposing Wnt-7a-positive protrusions was dependent on Wnt ligand palmitoleation (Figures 1L, O, and U) and calcium signalling (Figures 1R and U). We also confirmed that Wnt-7a-positive protrusions derived from post-synaptic neurites co-localise Wnt-7a and PSD95 and cluster of the pre-synaptic marker, Bassoon (BSN) in opposing neurites (Figures 1M and N). Furthermore, the co-localisation of Wnt-7a with PSD95 and co-clustering of Wnt-7a with BSN were also both dependent on Wnt palmitoleation (Figures 1P, Q, and U) and calcium signalling (Figures 1 S-U).

### Membrane-tethered Wnt phenocopies normal Wnt in clustering Wnt and synaptic components

The identification of Wnt-signalling filopodia markers and Wnt-7a on post-synaptic dendritic protrusions that induce clustering of LRP6 is suggestive of signalling *in trans* from the post-synaptic neuron to the pre-synaptic neuron. We wanted to identify if Wnt-7a could cause these signals without diffusion into the extracellular space. Therefore, we over-expressed a GFP-tagged active Wnt-7a in DIV60 iPSC-neurons and co-expressed morphotrap - the GFP nanobody Vhh-CD8-mCh^18^ - which tethers GFP-tagged proteins to the membrane through the single pass CD8 domain (Figures S2 and 2A). To ensure that Wnt-7a-GFP is efficiently trapped at the membrane, we co-transfected the Wnt ligand and morphotrap in a molar ratio of 1:1.5. Co-localisation analysis of dendritic filopodia showed a positive association of Wnt-7a-GFP with a membrane-associated mCherry (mem-mCh; Figure 2B). However, we found a significantly stronger association with morphotrap (Figures 2 C and D), suggesting efficient tethering of the Wnt-7a-GFP protein to the membrane. After co-transfecting Wnt-7a-GFP with either mem-mCh or morphotrap, we post-stained cultures for LRP6 after 24 h or PSD95 after 48 h. We found a significant induction of LRP6 clustering when Wnt-7a-GFP was over-expressed and a comparable significant induction if Wnt-7a-GFP was co-transfected with morphotrap (Figures 2E-H), suggesting that morpho-trapped Wnt-7a-GFP has a similar paracrine signalling activity compared to Wnt-7a-GFP. Furthermore, PSD95 expression on Wnt-7a-positive protrusions was significantly increased when Wnt-7a was over-expressed and similarly when it was membrane-tethered (Figures 2I-L). Our results suggest that membrane-tethered Wnt-7a – similar to Wnt-7a - can activate Wnt signalling *in trans* and promote synapse formation.

**Figure 2.**
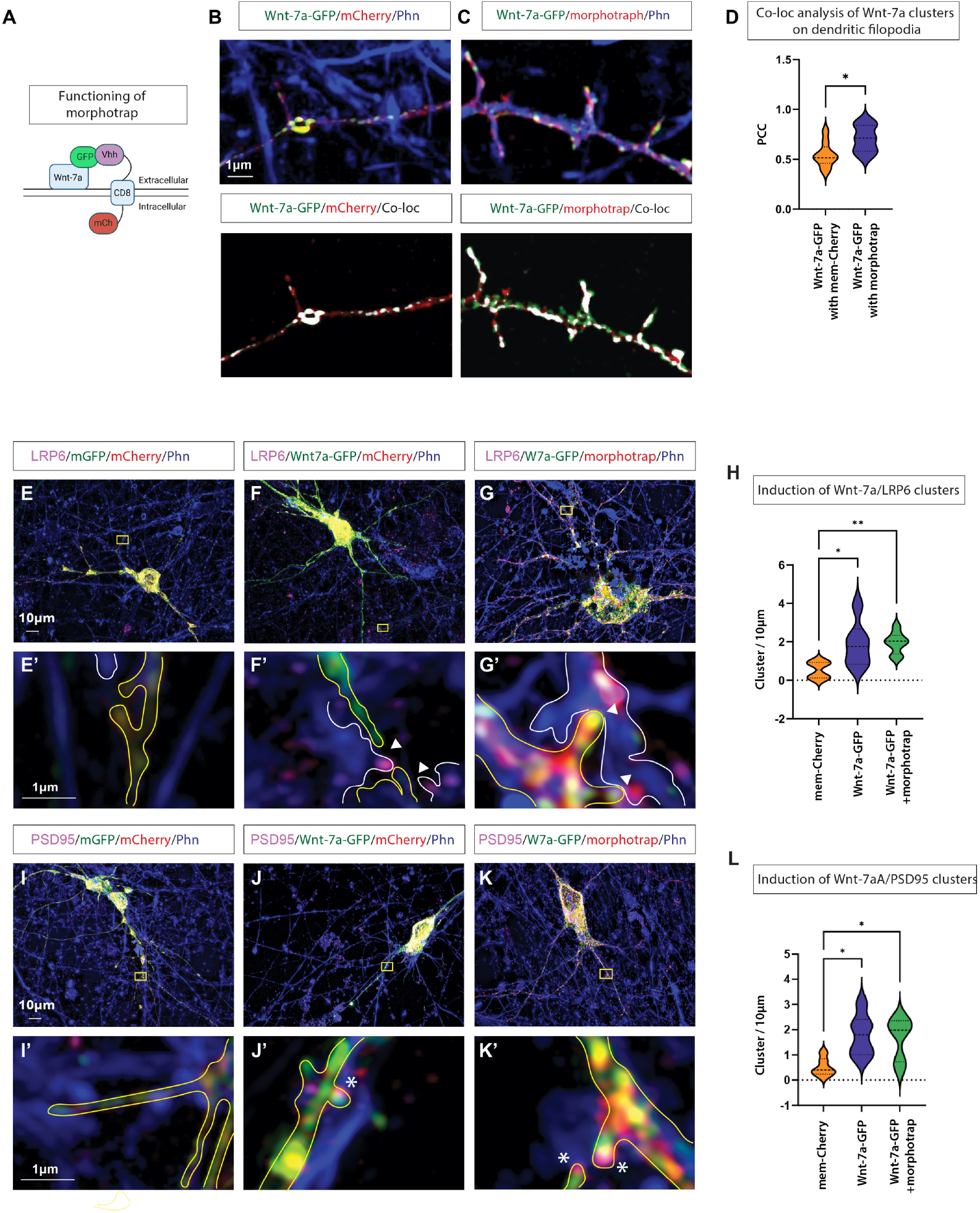
Membrane-tethered Wnt-7a-GFP can cluster Wnt and synaptic components. (**A**) Schematic representation of the tethering experiment using double transfection of Wnt-7a-GFP and the Morphotrap nanobody construct, Vhh-CD8-mCh that binds GFP-tagged proteins. (**B**) Colocalisation of mem-mCh and Wnt-7a-GFP on protrusions. (**C**) Significantly higher Wnt-7a-GFP colocalisation on filopodia is observed when co-transfected with Morphotrap, quantified by Pearson’s correlation coefficient (PCC; **D**). (**E, E’**) Transfection of iPSC-derived cortical neurons with mem-GFP and mem-mCh, followed by post-staining for LRP6 and the actin cytoskeleton (Phalloidin). (**F, F’**) Transfection of iPSC-derived cortical neurons with Wnt-7a-GFP and mem-mCh, followed by post-staining for LRP6 and the actin cytoskeleton (Phalloidin) identified Wnt-7a-GFP positive protrusions that cluster LRP6 in apposed cells. (**G, G’**) Transfection with Wnt-7a-GFP and Morphotrap, followed by post-staining for LRP6 and actin, identified protrusions harbouring membrane-tethered Wnt-7a-GFP that cluster LRP6 in apposed cells. (**H**) Quantification of co-clustering of LRP6 with Wnt-7a-GFP. (**I, I’**) Transfection of iPSC-derived cortical neurons with mem-GFP and mem-mCh, followed by post-staining for PSD95 and the actin cytoskeleton (Phalloidin). (**J, J’**) Transfection with Wnt-7a-GFP and mem-mCh, followed by post-staining for PSD95 and actin, found co-localisation on dendritic protrusions. (**K, K’**) Wnt-7a-GFP co-transfected with Morphotrap and co-localised with PSD95 on dendritic protrusions. (**L**) Quantification of co-localisation of PSD95 with Wnt-7a-GFP +/− morphotrap on dendritic protrusions. Statistical significance was addressed using One way ANOVA with Dunnett’s post hoc test for multiple comparisons, comparing groups to the mem-mCh/mem-GFP control group.

### Calcium transients on dendritic protrusions depend on membrane-associated Wnts

Next, we wanted to address the link between Wnt signal activation and calcium signalling at the dendritic filopodia contact sites because calcium signalling has been postulated to trigger the transformation of dendritic filopodia into functional spines of cortical neurons^19^. However, how calcium transients in filopodia guide precise synaptic formation needs to be better understood. Using live SIM imaging, we characterised filopodia contacts with an opposing neurite (Figures 3A and 3B) and found most connections were short-lived, with less than 15% forming stable contacts (defined as greater than 6 minutes; Figure 3C). Then, using a membrane-tethered calcium sensor (mem-gCAMP7S), we quantified the number of calcium transients in a 5 μm radius of the dendritic filopodia tips. We found increased calcium transients after stable connections had formed in the pre- and postsynaptic neurites (Figure 3D). Since we identified functional Wnt-7a-bearing cytonemes, we postulated that the combination of calcium signalling and localised Wnt signalling might provide a mechanism to guide dendritic filopodia to precise focal locations on the pre-synaptic neurite. To investigate this, we developed a membrane-bound ‘Wnt scissor’ to inactivate Wnt proteins specifically on the plasma membrane, here tips of dendritic filopodia (Figure S3). Therefore, we tethered Notum, a carboxylesterase that inactivates Wnt proteins by removing an essential palmitoleate moiety from Wnt proteins^20^, with a CD8 anchor to the membrane (memNotum; Figure S3A). We observed a strong localisation of memNotum to filopodia (Figure 3B), and the Wnt scissor memNotum efficiently removes Wnt-7a-GFP from the cytoneme tips (Figure S3C, D). Removal of membrane-associated Wnt-7a leads to a significant downregulation of SuperTOPFlash-based Wnt reporter activation in a co-culture setting (Figure 3 SE-I). With this tool in hand, we hypothesised that over-expressing memNotum preferentially target membrane-associated Wnts but leave extracellularly localised Wnts intact. We found that over-expression of memNotum significantly reduced calcium transients at filopodia contact sites (Figures 3F, H, I), supporting a role for membrane-associated Wnts in the initiation of synaptic formation from dendritic filopodia.

**Figure 3.**
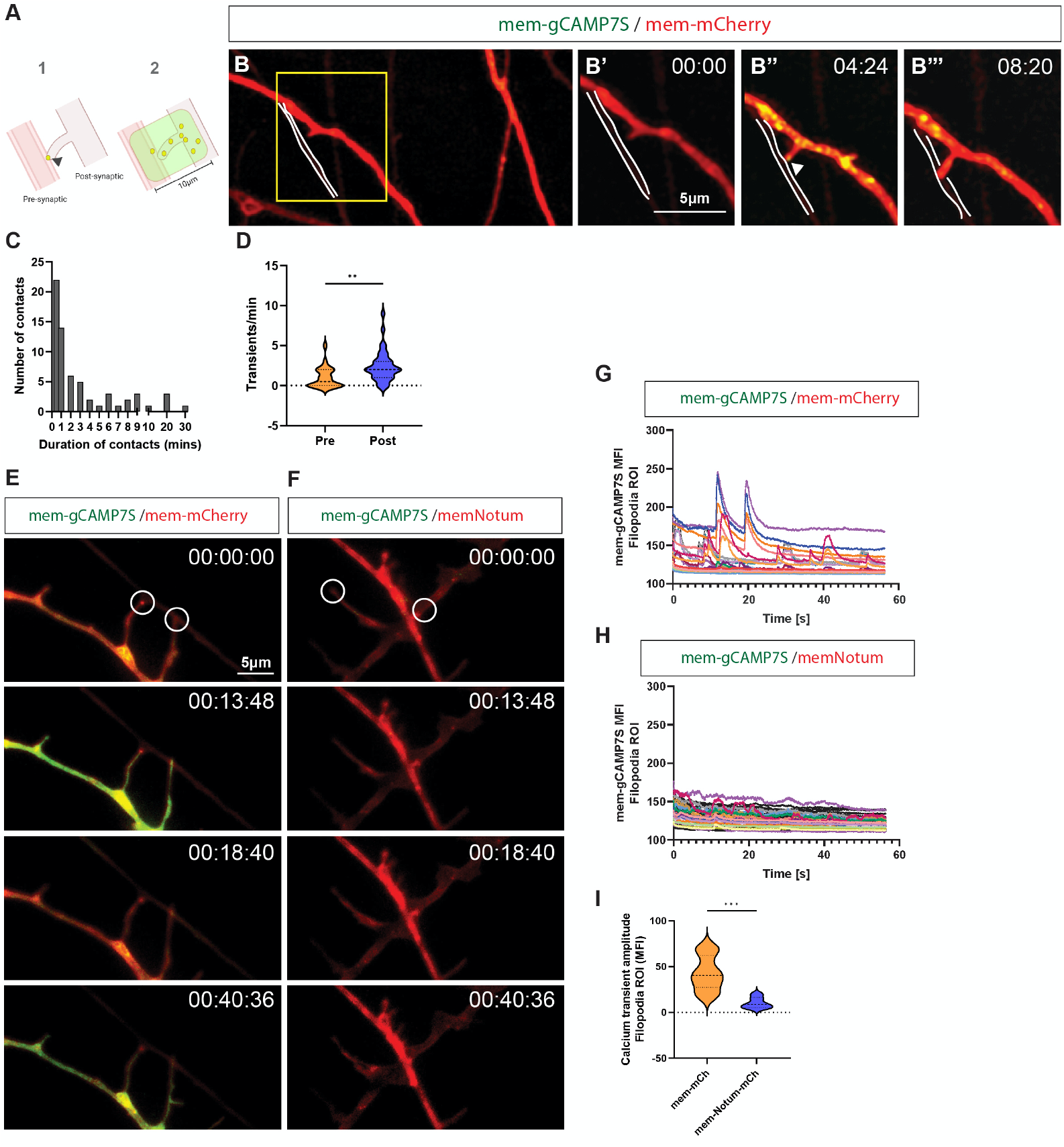
Calcium transients on dendritic protrusions depend on membrane-associated Wnts. (**A**) Schematic of calcium transient quantifications, with (1) showing the quantification of filopodia contacts with an oppositely apposed neurite and the duration of those contacts, and (2) showing the quantification of the number of transients occurring pre-contact vs post-stabilisation within a 5μm radius of the contact point. (**B**) Live imaging of iPSC-derived cortical neurons transfected with a membrane marker (mem-mScarlet-I) and a membrane-tethered calcium sensor (mem-gCAMP7S). Cells were imaged continuously for 10 seconds with 1-minute intervals using Apotome SIM. (**B’’**) Upon initial contact with another neurite, calcium transients are seen to localise at the tips of dendritic filopodia (white arrowhead). (**B’’’**) Upon stable contact, transients occur more frequently close to the newly formed connection. (**C**) Quantification of contact number and duration was performed on a subset of experiments (10 fields of view from two biological replicates) and shows most calcium-associated filopodial contacts are transient, with only a small proportion (<15%) stable for over 6 minutes. (**D**) Quantifying the number of calcium transients occurring within a 5μm radius of the dendritic filopodia tip shows an increase in transients after forming a stable connection. The experiment was performed in biological triplicate, with at least 8 fields of view (20μm^2^) analysed/repeated. Statistical significance was addressed using Student’s t-test to compare the number of transients/min pre-stable contact and transients/min post-stable contact. Stable contacts were defined by being present >6 minutes of the experimental recording time, **P < .01. (**E**) Continuous calcium imaging (1 minute bursts) was performed using mem-gCAMP7S on neurons co-transfected with mem-mCh in HILO mode to obtain 10μm optical sections (mm:ss: ms). (**F**) Continuous calcium imaging was performed on neurons co-transfected with the Wnt ‘scissor’ memNotum, and mem-gCAMP7S (mm:ss:ms). (**G**) Traces of membrane-associated calcium transients at protrusion tips in control neurons. (**H**) Traces of membrane-associated calcium transients at protrusion tips in memNotum-transfected neurons. (**I**) Quantifying calcium transient amplitude at dendritic protrusion tips identified a significant reduction in neurons transfected with memNotum. Statistical significance was addressed using Student’s t-test, ***P < 0.001.

### Membrane-tethered Wnts are required for synaptogenesis

Finally, we wanted to identify if the inactivation of membrane-associated Wnts was sufficient to reduce the synaptogenic capacity of dendritic filopodia. Therefore, we transfected iPSC-derived cortical neurons with a *Fibronectin intrabodies generated by mRNA display* (*FingRs*) construct that tags endogenous levels of PSD95 with GFP (FingR-PSD95)^21^. FingR -PSD95 transfection of control neurons showed PSD95 puncta localising to dendritic protrusions (Figure 4A). Next, we inhibited the function of membrane-associated Wnts by memNotum and observed that the neurons co-transfected with FingR -PSD95 and memNotum display significantly reduced PSD95-positive protrusions (Figures 4B and F). Similarly, soluble Notum treatment significantly reduced PSD95-positive protrusions, suggesting that most of the Wnts are membrane-associated during synaptogenesis (Figures 4C and F). LP-922056 treatment of memNotum transfected neurons reversed the PSD95-positive protrusion reduction (Figure 4D and F), suggesting a specific function of memNotum. Interestingly, transfection of neurons with memNotum, treatment of control cells with soluble Notum, or treatment of memNotum transfected neurons with the Notum inhibitor, LP-922056, did not affect protrusion number (Figures 4E), suggesting that the presence of dendritic filopodia without membrane-localised Wnt is not sufficient for PSD95 clustering. Finally, we wanted to address if membrane-tethered Wnts are also required for clustering of pre-synaptic markers, because co-localisation of the post-synaptic compartment with the pre-synaptic active zone is required to confirm synapse formation. Therefore, we performed a similar experiment as outlined above, but post-stained cultures with antibodies against post-synaptic PSD95 and pre-synaptic BSN and performed confocal microscopy and subsequent quantification of the number of PSD95/BSN clusters localised at the protrusions of transfected neurons (Figures 4G-K). We found a significant reduction in PSD95/BSN puncta when neurons were transfected with memNotum (Figures 4H) or treated with soluble Notum (Figure 4I). The memNotum-induced decline in PSD95/BSN clustering could be reversed with treatment with the Notum inhibitor LP-922056 (Figure 4J).

**Figure 4.**
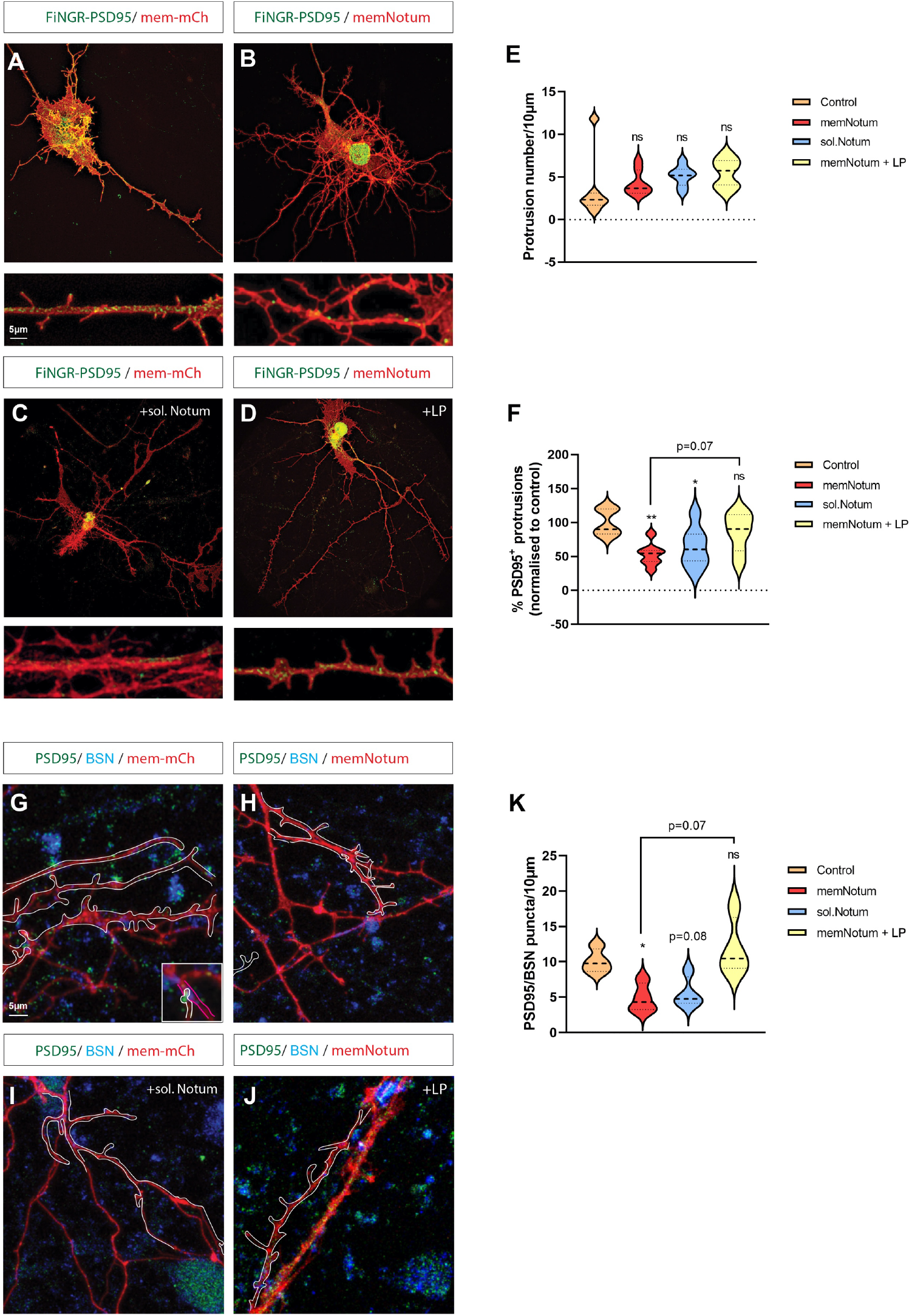
Membrane-tethered Wnts are required for synaptogenesis. Cortical neurons were transfected with FiNGR-PSD95-GFP to tag PSD95 endogenously. Control neurons were co-transfected with mem-mCh and showed strong PSD95 puncta localisation to protrusions (**A**), which is lost when neurons were co-transfected with memNotum (**B**) or when control cultures were incubated with soluble, recombinant Notum (**C**). Puncta expression could be rescued in memNotum transfected neurons by pre-treatment with Notum inhibitor LP-922056 (**D**). (**E**) Quantification of protrusion number showed no difference between treatment groups, (**F**) PSD95-positive protrusion quantification. Neurons were stained for PSD95 and Bassoon (BSN) after transfection with either mem-mCh (**G**) or memNotum (**H**), and co-localised puncta in mCh-expressing protrusions were quantified (**K**). PSD95/BSN puncta (shown in the inset in **G**) were significantly reduced in memNotum-transfected neurons compared to control, but not when control cultures were incubated with soluble, recombinant Notum (**I**). Puncta expression could be rescued in memNotum transfected neurons by pre-treatment with LP-922056 (**J**). Statistical significance was addressed using one-way ANOVA with Dunnett’s multiple comparison test to compare relevant controls within groups and Student’s t-test to compare specific combinations. *P < 0.05; **P < 0.01; ***P < 0.005; ****P < 0.001. Experiments were performed in biological triplicate, with at least 5 fields/group were analysed/replicated.

## Discussion

Wnt7a signals through the Wnt/β-catenin pathway to regulate the formation and maturation of presynaptic and postsynaptic structures and is necessary for normal synaptic function^22^. For example, Wnt7a/β-catenin signalling has been shown to promote spine formation and PSD95 expression in the murine hippocampus^23^. Wnt7a has also been shown to increase calcium entry to activate CaMKII in spines leading to a growth in excitatory synapse formation in mice^10^. Although the beneficial role of Wnt7a has been studied, it is still unclear how Wnt7a is transported between the post- and presynaptic membrane to allow the establishment of a functional spine at a precise location.

By using super-resolution LSIM, we show that cytoneme-like structures, here dendritic filopodia, are decorated with Wnt-7a and transport the ligand to neighbouring neurons to activate Wnt signalling locally. We show that Wnt-7a subsequently activates calcium transients and clustering of pre- and post-synaptic markers at the contact site of the dendritic filopodium, suggesting a requirement for membrane association of Wnts during synaptogenesis. In support of our hypothesis, data from *Drosophila* suggest that a membrane-tethered Wnt/Wg can function equally compared to a WT-WNT/Wg in wing development^24^. Thus, our data suggests that lipophilic Wnt proteins are not freely diffusible but maintain membrane association. Long-range signalling might require a specific transport mechanism, such as an exovesicle-based transport or the transport on cytonemes^25^. We provide the first evidence that dendritic filopodia are Wnt-7a transporting cytonemes. In support of our finding, accumulating evidence suggests that these thin and highly polarised cytonemes can also operate in other contexts, such as during the transport of Wnt8a proteins in zebrafish gastrulation^26,27^, Wnt2b in the mouse intestinal crypt^12^, Wnt3a in human embryonic stem cells^28^, and Wnt3 in gastric cancer^13^.

Can it be excluded that a meaningful proportion of the Wnt-7a protein is also freely diffusible? In our experiments, we find no evidence for the requirement of an additional pool of Wnt-7a other than the signals transported on dendritic filopodia. For example, we show that membrane-tethered Wnt-7a is similar in function to normal Wnt-7a, suggesting a continuous association with membranes during synaptogenesis. Furthermore, the inactivation of membrane-tethered Wnt proteins on the post-synaptic neuron cell is sufficient to inhibit the induction of calcium transients and synaptic protein clustering. Moreover, WT neurons in the same cell culture dish cannot rescue the lack of membrane-bound Wnt-7a on dendritic filopodia tips needed for synapse formation, arguing against a pool of freely diffusible Wnt-7a or ligand-loaded exovesicles.

In conclusion, our work bridges a fundamental knowledge gap and provides an explanation of how Wnt-7a functions in excitatory synapse formation. Dendritic filopodia are necessary for mobilising Wnt-7a and trafficking the ligand to dendrites of neighbouring neurons. Then, Wnt-7a on the tips of dendritic cytonemes triggers the local induction of calcium signalling leading to the clustering of pre- and post-synaptic markers and the formation of new, functional synapses.

## Acknowledgements

We thank K. Fang for cell culture maintenance and construct cloning. We thank C. Liddle and the Bioimaging Facility for training and support in super-resolution microscopy and maintenance of the Zeiss Elyra7 Lattice SIM system. We thank J.P. Vincent for the kind gift of the hNotumFL construct. This work was supported by funding through BRACE Dementia Research (S.S., A.B. and R.K.), a Wellcome Trust Institutional Strategic Support Award (TREE) award to T.M.P and S.S. (Grant number 204909/Z/16/Z), a BBSRC equipment grant (19ALERT BB/T017899/1, awarded to SS et al.), and the Living Systems Institute.

## Materials and Methods

### Plasmids and antibodies

The following plasmids were used in transfections: pCS2+-membrane-mCherry (Mattes et al., 2018), pCAG-mGFP membrane-bound GFP (Addgene 14757), pcDNA3.1-Wnt7a-GFP, 7×TRE-SuperTOPFlash-NLS-mCherry^29^, pCS2+-LifeAct-GFP, pCAG-PSD95.FingR-eGFP-CCR5TC (Addgene 46295), Lck-mScarlet-I (Addgene 98821), pCS2+-Vhh-CD8-mCherry (cloned from Gal4/LexA-Vhh-CD8-mCherry (Affolter lab) into pCS2+ vector using XhoI/XbaI), pCS2+-GAP43-jGCaMP7s (cloned from pGP-CMV-jGCaMP7s (Addgene 104463) into pCS2+-GAP43-GFP using XbaI/SnaBI), pcDNA3.1-Notum-CD8-mCh (assembled by Gibson cloning from pcDNA3.1-hNotumFL (Gift from J.P Vincent) and pCS2+-Vhh-CD8-mCherry).

The following primary antibodies were used for immunofluorescence: anti-Wnt7a (abcam; ab100792), anti-LRP6 (BioTechne; FAB1505R), anti-Bassoon (abcam; ab82958), anti-PSD95 (abcam; ab18258), anti-PSD95 (abcam; ab13552), anti-Flotillin-2 (Santa Cruz; sc-28320), anti-Flotillin-2 (abcam; ab113661), anti-Myosin-X (Santa Cruz C-1; sc-166720), anti-Evi (EMD Milipore; YJ5). The following AlexaFluor (Thermofisher) secondary antibodies were used for immunofluorescence: goat anti-rabbit 488 (abcam; ab150077), goat anti-mouse 647 (abcam; ab150115), donkey anti-mouse 488 (abcam; ab150105) donkey anti-goat 647 (abcam; ab150135).

### Cell culture

#### iPSC maintenance

iPSCs (GM23280A, KOLF2.1J) were obtained from the Coriell Institute for Medical Research and the iNDI consortium (gift from Prof W. Skarnes), respectively. Lines were maintained as colonies on human ES-qualified Matrigel (Corning) in StemFlex (StemCell Technologies). Colonies were routinely passaged in a 1:6 split using EDTA and banked. All cell lines were tested regularly for mycoplasma by endpoint PCR testing every 3 months and broth tests every 12 months.

### Differentiation of iPSCs into cortical neurons

Neurons were derived as previously described^30^. Briefly, iPSCs were plated as colonies onto Matrigel and differentiated by treatment with neuronal differentiation media (DMEM/F12:Neurobasal in a 1:1 ratio, HEPES 10mM, N2 supplement 1%, B27 supplement 1%, Glutamax 1%, ascorbic acid 5uM, insulin 20ug/ml) supplemented with SB431542 (10uM) and LDN8312 (0.2μM) from D0-D12, replacing media daily. On D12, cultures were replated using accutase in neuronal differentiation media supplemented with bFGF (20ng/ml), CHIR-99021 (1μM), and Y-27632 (50μM). On D13-D18, media was changed daily (neuronal differentiation media supplemented with bFGF (20ng/ml), CHIR-99021 (1μM). Cells were either banked as neural progenitor cells (NPCs), stained for cortical neuron lineage markers, or maintained in culture until D60-80 for experimentation. Cells were maintained in differentiation media supplemented with L-Ascorbic acid (200μM), BNDF (20ng/ml), GDNF (10ng/ml), and Compound E (0.1μM) for 7 days before Compound E was removed. Cultures were replenished every 3-4 days with a 50:50 media change. Treatments with 1μg/ml of recombinant human NOTUM protein (Bio-Techne) or 100nM LP-922056 (Kind gift from ARUK DDI) were performed for 48 h prior image acquisition.

### Neuronal transfection

Neuronal cultures were transfected with equimolar ratios of plasmid DNA totalling 1.5 μg, unless otherwise stated, using a calcium-phosphate method as previously described^31^. A plasmid/CaCl_2_ (12.4mM) master mix was prepared for each combination to be transfected in Hank’s balanced salt solution (HBSS), mixing gradually (1/8 of DNA:CaCl_2_ mixture/addition) to prevent the formation of large transfection complexes. Complexes were incubated for 20 minutes at RT, then added to cultures (no more than 1/10 of total volume) and incubated at 37°C for 4.5 h, before a sodium acetate (300mM in DMEM:F12) wash at 37°C for 5 minutes to dissolve precipitated complexes. Cultures were maintained in neuronal differentiation media supplemented for 24-48 h prior to imaging analyses.

### AGS cell culture and transfection

The primary gastric adenocarcinoma cell line AGS was a kind gift from Dr Toby Phesse (Cardiff University, UK). AGS cells were maintained in RPMI-1640 (Sigma-Aldrich), supplemented with 10% FBS. In addition, cells were routinely passaged with TrypLE (Thermofisher). Transient transfections of AGS cells were performed using FuGeneHD (Promega) according to the manufacturer’s protocol (using a 3:1 FuGene: DNA ratio).

### SuperTOPFlash-based Wnt reporter assay

To determine the functionality of the membrane-tethered Notum construct, 0.5×10^6^ AGS cells were reverse transfected with SuperTOPFlash reporter plasmid (7×TRE-NLS-mCherry), and 0.5×10^6^ AGS cells reverse transfected with indicated plasmids in 6 well plates. After 24 hr incubation, both cell types were trypsinised, counted, and 0.5×10^6^ of each population were co-cultivated in 6 well plates for a further 48 hr before fixation and post-staining with DAPI for total nuclei quantification. Image acquisition was performed on a Leica DMI6000 SD using a 20x objective. For all assays, at least 4 images were taken in random locations for each biological repeat. The fluorescence intensity of mCherry-positive nuclei was measured on Fiji software. The number of DAPI-positive nuclei was also counted as a measure of proliferation.

### Antibody staining and image acquisition

Cells were plated onto 1.5H precision coverslips and following indicated treatment/incubation cells were immediately fixed using modified MEM-Fix (4% formaldehyde, 0.25–0.5% glutaraldehyde, 0.1 M Sorenson’s phosphate buffer, pH7.4)^32,33^ for 7 min at 4°C. Aldehydes were subsequently quenched by incubation with NaBH_4_ (0.1% w/v) for 7 minutes at RT. Further quenching was performed by 3 x 10 min washes in PBS-Glycine (0.2M). Cells were then incubated in permeabilisation solution (0.1% Triton-X-100, 5% serum, 0.1 M glycine in 1×PBS) for 1 hr at RT. Primary antibodies were diluted in incubation buffer (0.1% Tween20, 5% serum in 1×PBS) and coverslips mounted on 40 μl spots overnight at 4°C in a humid environment. Coverslips were then washed with 1x PBS 3 x for 5 min before mounting on 40 μl spots of secondary antibodies ± Phalloidin-405 (for actin cytoskeleton experiments) diluted in incubation buffer for 45 minutes at RT. Coverslips were then washed 3 x for 5 min with 1x PBS before mounting onto glass slides using ProLong Diamond Antifade mountant (Invitrogen) and left to dry for 24 hr at 4°C before imaging. Confocal microscopy was performed on an inverted Leica TCS SP8 X laser-scanning microscope using the 63x water objective. Structured Illumination Microscopy for transfection studies, immunofluorescent antibody staining, and live cell imaging were performed on the Zeiss Elyra7 Lattice SIM system using 40x and 63x objectives, employing Apotome and Lattice SIM modes, respectively. In addition, live calcium imaging was performed on the Elyra7 Lattice SIM system in laser widefield, HILO mode with cells cultured in glass bottom dishes (Ibidi) in Image Optimised Medium (Stem Cell Technologies).

### Quantifications and statistical analyses

All protrusion quantifications were calculated from Z-stack images of cells stained with Phalloidin or expressing membrane-mCherry, or lck-mScarlet-I using Imaris software (Oxford Instruments). Protrusion length was measured from the tip of the filopodia to the base, where it contacted the neurite. In the case of branching protrusions, one branch (the longest) would be measured. Colocalisation quantifications were performed using the ‘Coloc’ function in Imaris using a protrusion tip threshold and normalised/10μm length of the neurite. All experiments/conditions were repeated in biological triplicates. At least 500μm of dendritic arborisation was analysed/experiment Significance was tested using Student’s t-test (parametric) against the relative controls. One-way ANOVA with Dunnett’s post hoc test was performed for multiple comparisons.

## Supplementary data

**Supplementary Figure 1. related to Figure 1.**
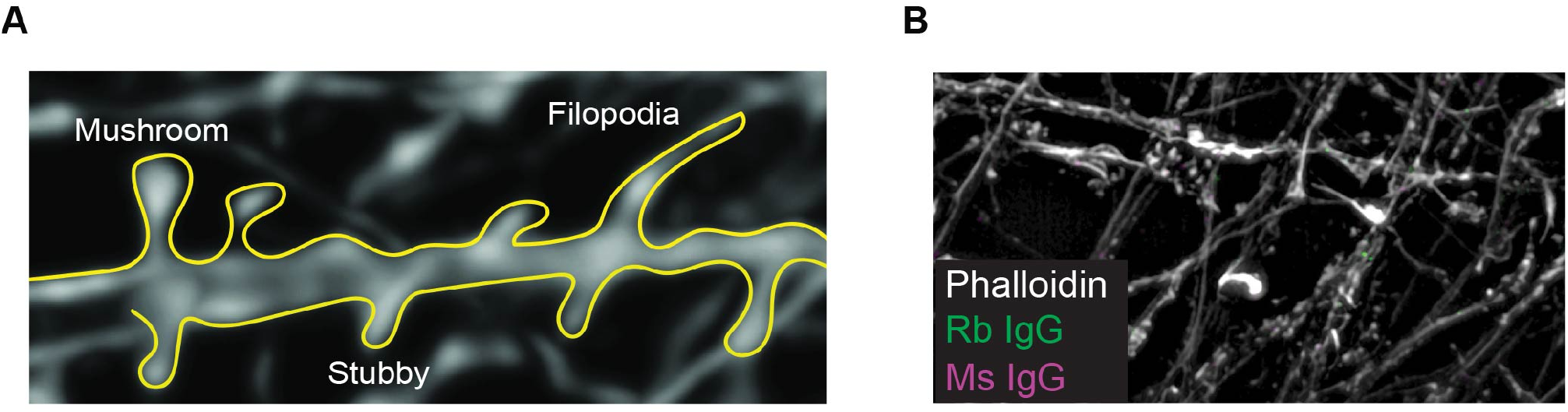
Quantification methodology and negative controls. (**A**) Outline of morphological characteristics used in quantification of dendritic protrusions. (**B**) IgG isotype controls for primary antibodies.

**Supplementary Figure 2. related to Figure 2.**
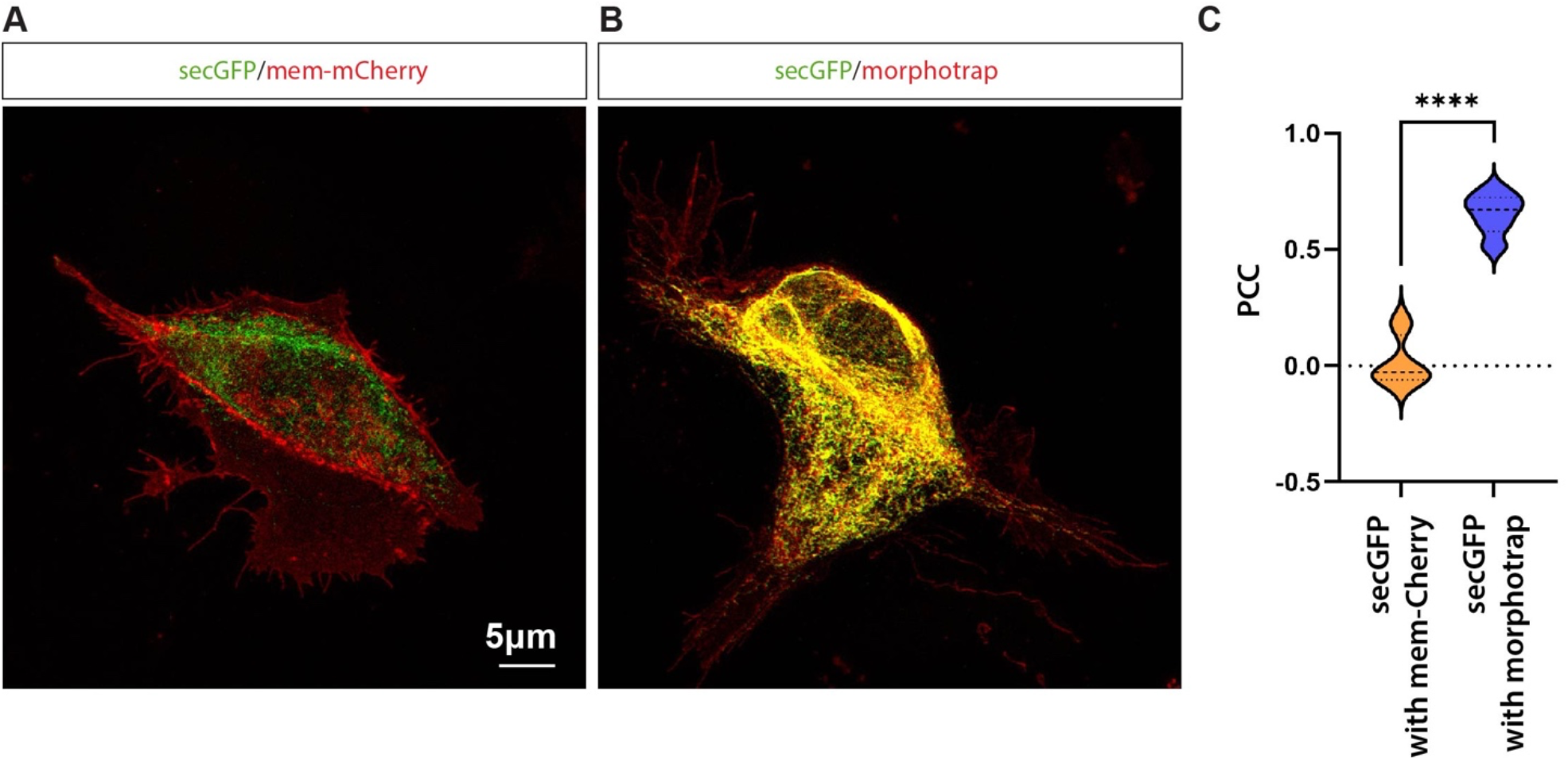
Morphotrap characterisation. (**A**) AGS cells transfected with soluble-GFP (secGFP) and mem-mCh. (**B**) AGS cells transfected with secGFP and Morphotrap. (**C**) Quantification identifies a strong colocalisation between the normally secreted soluble GFP (secGFP) and the Morphotrap.

**Supplementary Figure 3, related to Figure 3.**
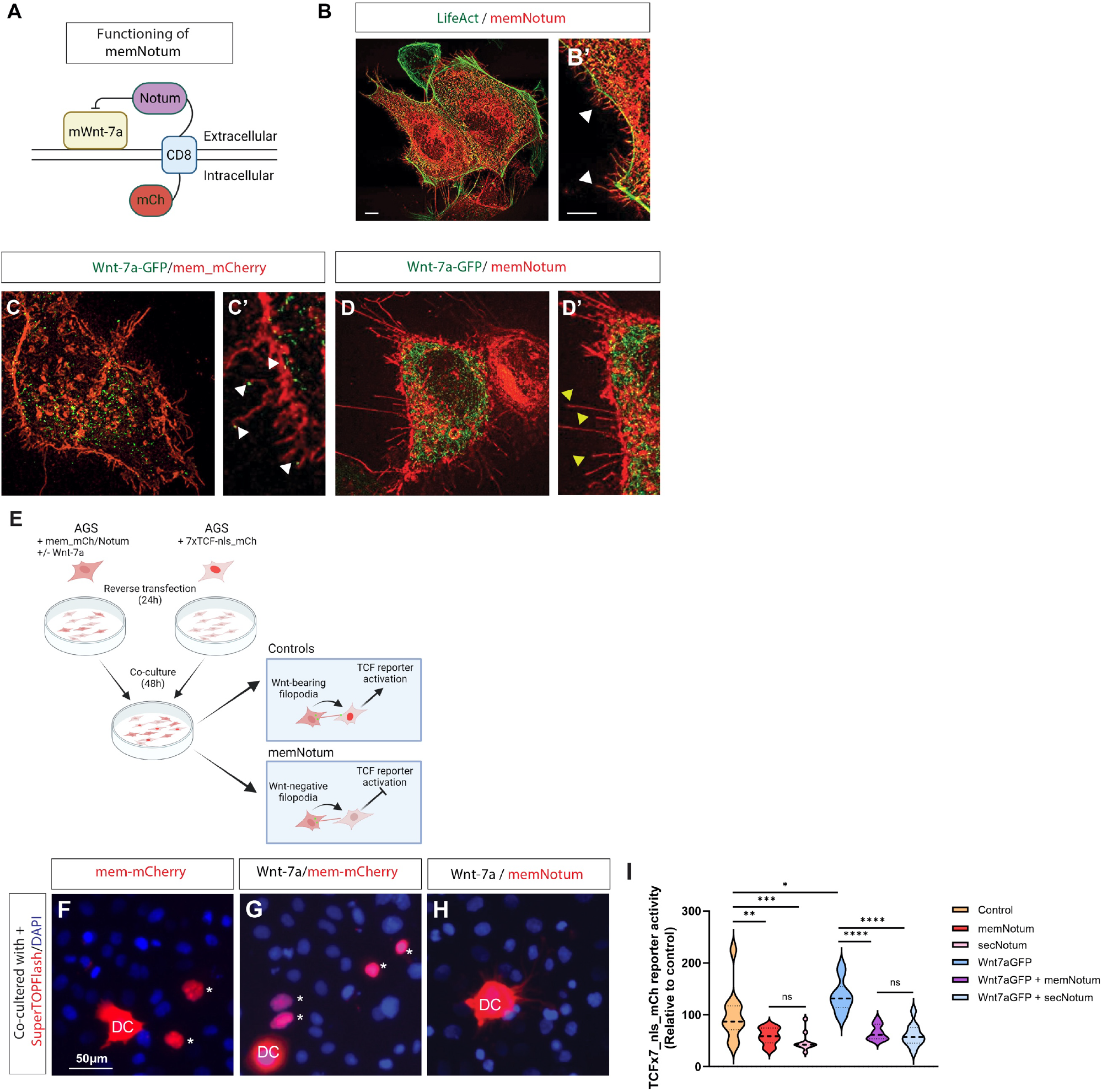
Generation and characterisation of membrane-tethered Wnt ‘scissor’. (**A**) Schematic representation of the generated mCherry-tagged membrane-tethered Notum construct (memNotum). (**B**) Transfection of human AGS cells with memNotum shows strong expression on all cell membranes and cellular filopodia (**B’**). Cells were co-transfected with LifeAct-GFP for visualisation of the actin cytoskeleton. (**C**) Co-transfection of the membrane marker, mem-mCh, and Wnt-7a-GFP show localisation of Wnt-7a-GFP to filopodia tips (**C’**; white arrowheads). (**D**) Co-transfection of memNotum and Wnt-7a-GFP reduces Wnt-7a-GFP positive filopodia (**D’**; yellow arrowheads). (**E**) Schematic of SuperTOPFlash-based TCFx7-nls-mCh reporter assay set-up. (**F**) TCF reporter expression can be observed in cells around the control donor cell (DC) population (mem-Ch; asterisks). (**G**) Reporter activity is increased if the donor cell population is co-transfected with Wnt-7a-GFP (mem-mCh; asterisks). (**H**) TCF reporter activity is significantly reduced if donor cells are transfected with memNotum. (**I**) Quantifying mCherry nuclear expression in the receiving cell population identified the ability of memNotum to significantly reduce Wnt-mediated paracrine signalling to a similar level to secreted Notum (transfection of full-length untethered human Notum. Statistical significance was addressed using one-way ANOVA with Dunnett’s multiple comparison test to compare relevant controls within groups and Student’s t-test to compare specific combinations. *P < .05; **P < .01; ***P < .005; ****P < .001. Experiments were performed in biological triplicate, with at least 3 fields/group were analysed/repeated.

## References

1. Ziv, N.E., and Smith, S.J. (1996). Evidence for a Role of Dendritic Filopodia in Synaptogenesis and Spine Formation. Neuron 17, 91–102. 10.1016/S0896-6273(00)80283-4.

2. Südhof, T.C. (2017). Synaptic Neurexin Complexes: A Molecular Code for the Logic of Neural Circuits. Cell 171, 745–769. 10.1016/J.CELL.2017.10.024.

3. Biederer, T., Sara, Y., Mozhayeva, M., Atasoy, D., Liu, X., Kavalali, E.T., and Südhof, T.C. (2002). SynCAM, a synaptic adhesion molecule that drives synapse assembly. Science 297, 1525–1531. 10.1126/SCIENCE.1072356.

4. Packard, M., Mathew, D., and Budnik, V. (2003). WNTS AND TGFβ IN SYNAPTOGENESIS: OLD FRIENDS SIGNALLING AT NEW PLACES. Nat Rev Neurosci 4, 113. 10.1038/NRN1036.

5. Qi, C., Luo, L. da, Feng, I., and Ma, S. (2022). Molecular mechanisms of synaptogenesis. Front Synaptic Neurosci 14, 55. 10.3389/FNSYN.2022.939793/BIBTEX.

6. McLeod, F., and Salinas, P.C. (2018). Wnt proteins as modulators of synaptic plasticity. Curr Opin Neurobiol 53, 90–95. 10.1016/J.CONB.2018.06.003.

7. Lai, S.L., Chien, A.J., and Moon, R.T. (2009). Wnt/Fz signaling and the cytoskeleton: potential roles in tumorigenesis. Cell Research 2009 19:5 19, 532–545. 10.1038/cr.2009.41.

8. Routledge, D., and Scholpp, S. (2019). Mechanisms of intercellular Wnt transport. Development 146. 10.1242/DEV.176073.

9. Hall, A.C., Lucas, F.R., and Salinas, P.C. (2000). Axonal remodeling and synaptic differentiation in the cerebellum is regulated by WNT-7a signaling. Cell 100, 525–535. 10.1016/S0092-8674(00)80689-3.

10. Ciani, L., Boyle, K.A., Dickins, E., Sahores, M., Anane, D., Lopes, D.M., Gibb, A.J., and Salinas, P.C. (2011). Wnt7a signaling promotes dendritic spine growth and synaptic strength through Ca 2+/Calmodulin-dependent protein kinase II. Proc Natl Acad Sci U S A 108, 10732–10737. 10.1073/PNAS.1018132108/-/DCSUPPLEMENTAL.

11. Dickins, E.M., and Salinas, P.C. (2013). Wnts in action: from synapse formation to synaptic maintenance. Front Cell Neurosci 7. 10.3389/FNCEL.2013.00162.

12. Mattes, B., Dang, Y., Greicius, G., Kaufmann, L.T., Prunsche, B., Rosenbauer, J., Stegmaier, J., Mikut, R., Özbek, S., Nienhaus, G.U., et al. (2018). Wnt/PCP controls spreading of Wnt/β-catenin signals by cytonemes in vertebrates. Elife 7. 10.7554/ELIFE.36953.

13. Routledge, D., Rogers, S., Ono, Y., Brunt, L., Meniel, V., Tornillo, G., Ashktorab, H., Phesse, T., and Scholpp, S. (2022). The scaffolding protein Flot2 promotes cytoneme-based transport of Wnt3 in gastric cancer. Elife 11. 10.7554/ELIFE.77376.

14. Du, L., Sohr, A., Li, Y., and Roy, S. (2022). GPI-anchored FGF directs cytoneme-mediated bidirectional contacts to regulate its tissue-specific dispersion. Nat Commun 13. 10.1038/S41467-022-30417-1.

15. Kornberg, T.B., and Roy, S. (2014). Cytonemes as specialized signaling filopodia. Development (Cambridge) 141, 729–736. 10.1242/DEV.086223.

16. Huang, H., Liu, S., and Kornberg, T.B. (2019). Glutamate signaling at cytoneme synapses. Science (1979) 363, 948–955. 10.1126/SCIENCE.AAT5053/FORMAT/PDF.

17. Bischoff, M., Gradilla, A.C., Seijo, I., Andrés, G., Rodríguez-Navas, C., González-Méndez, L., and Guerrero, I. (2013). Cytonemes are required for the establishment of a normal Hedgehog morphogen gradient in Drosophila epithelia. Nat Cell Biol 15, 1269–1281. 10.1038/NCB2856.

18. Harmansa, S., Hamaratoglu, F., Affolter, M., and Caussinus, E. (2015). Dpp spreading is required for medial but not for lateral wing disc growth. Nature 2015 527:7578 527, 317–322. 10.1038/nature15712.

19. Lohmann, C., and Bonhoeffer, T. (2008). A Role for Local Calcium Signaling in Rapid Synaptic Partner Selection by Dendritic Filopodia. Neuron 59, 253–260. 10.1016/J.NEURON.2008.05.025.

20. Kakugawa, S., Langton, P.F., Zebisch, M., Howell, S.A., Chang, T.H., Liu, Y., Feizi, T., Bineva, G., O’Reilly, N., Snijders, A.P., et al. (2015). Notum deacylates Wnt proteins to suppress signalling activity. Nature 2015 519:7542 519, 187–192. 10.1038/nature14259.

21. Gross, G.G., Junge, J.A., Mora, R.J., Kwon, H.B., Olson, C.A., Takahashi, T.T., Liman, E.R., Ellis-Davies, G.C.R., McGee, A.W., Sabatini, B.L., et al. (2013). Recombinant Probes for Visualizing Endogenous Synaptic Proteins in Living Neurons. Neuron 78, 971–985. 10.1016/J.NEURON.2013.04.017.

22. Teo, S., and Salinas, P.C. (2021). Wnt-Frizzled Signaling Regulates Activity-Mediated Synapse Formation. Front Mol Neurosci 14. 10.3389/FNMOL.2021.683035.

23. Ramos-Fernández, E., Tapia-Rojas, C., Ramírez, V.T., and Inestrosa, N.C. (2019). Wnt-7a Stimulates Dendritic Spine Morphogenesis and PSD-95 Expression Through Canonical Signaling. Mol Neurobiol 56, 1870–1882. 10.1007/S12035-018-1162-1.

24. Alexandre, C., Baena-Lopez, A., and Vincent, J.P. (2014). Patterning and growth control by membrane-tethered Wingless. Nature 505, 180–185. 10.1038/NATURE12879.

25. Zhang, C., and Scholpp, S. (2019). Cytonemes in development. Curr Opin Genet Dev 57, 25. 10.1016/J.GDE.2019.06.005.

26. Stanganello, E., Hagemann, A.I.H., Mattes, B., Sinner, C., Meyen, D., Weber, S., Schug, A., Raz, E., and Scholpp, S. (2015). Filopodia-based Wnt transport during vertebrate tissue patterning. Nature Communications 2015 6:1 6, 1–14. 10.1038/ncomms6846.

27. Brunt, L., Greicius, G., Rogers, S., Evans, B.D., Virshup, D.M., Wedgwood, K.C.A., and Scholpp, S. (2021). Vangl2 promotes the formation of long cytonemes to enable distant Wnt/β-catenin signaling. Nature Communications 2021 12:1 12, 1–19. 10.1038/s41467-021-22393-9.

28. Junyent, S., Garcin, C.L., Szczerkowski, J.L.A., Trieu, T.J., Reeves, J., and Habib, S.J. (2020). Specialized cytonemes induce self-organization of stem cells. Proc Natl Acad Sci U S A 117, 7236–7244. 10.1073/PNAS.1920837117/-/DCSUPPLEMENTAL.

29. Moro, E., Ozhan-Kizil, G., Mongera, A., Beis, D., Wierzbicki, C., Young, R.M., Bournele, D., Domenichini, A., Valdivia, L.E., Lum, L., et al. (2012). In vivo Wnt signaling tracing through a transgenic biosensor fish reveals novel activity domains. Dev Biol 366, 327–340. 10.1016/J.YDBIO.2012.03.023.

30. Strano, A., Tuck, E., Stubbs, V.E., and Livesey, F.J. (2020). Variable Outcomes in Neural Differentiation of Human PSCs Arise from Intrinsic Differences in Developmental Signaling Pathways. Cell Rep 31, 107732–107732. 10.1016/J.CELREP.2020.107732.

31. Hawkins, S., Namboori, S.C., Tariq, A., Blaker, C., Flaxman, C., Dey, N.S., Henley, P., Randall, A., Rosa, A., Stanton, L.W., et al. (2022). Upregulation of β-catenin due to loss of miR-139 contributes to motor neuron death in amyotrophic lateral sclerosis. Stem Cell Reports 17, 1650–1665. 10.1016/J.STEMCR.2022.05.019.

32. Bodeen, W.J., Marada, S., Truong, A., and Ogden, S.K. (2017). A fixation method to preserve cultured cell cytonemes facilitates mechanistic interrogation of morphogen transport. Development 144, 3612–3624. 10.1242/DEV.152736.

33. Rogers, S., and Scholpp, S. (2020). Preserving Cytonemes for Immunocytochemistry of Cultured Adherent Cells. Methods in Molecular Biology 2346, 183–190. 10.1007/7651_2020_305.

